# Carbonic anhydrase plays multiple roles in acetotrophic growth of a model marine methanogen from the domain *Archaea*

**DOI:** 10.1101/2024.04.24.590269

**Authors:** Michel Geovanni Santiago-Martínez, Sabrina Zimmerman, Ethel E. Apolinario, Kevin R. Sowers, James G. Ferry

**Affiliations:** Department of Biochemistry and Molecular Biology, Pennsylvania State University, University Park, Pennsylvania, 16801, USA; Department of Marine Biotechnology, Institute of Marine and Environmental Technology, University of Maryland, Baltimore County, Baltimore, Maryland, 21202, USA

**Keywords:** archaea, global warming, carbon cycle, anaerobic, methane

## Abstract

Carbonic anhydrase (CA) catalyzes the reversible hydration of CO_2_ to bicarbonate and a proton. The enzyme is universally distributed in all three domains of life and plays diverse physiological roles in the domains *Eukarya* and *Bacteria*. Remarkably, a physiological role has not been identified for any CA from the domain *Archaea*. Herein are described roles for a gamma class CA (Cam) from the methane-producing marine archaeon *Methanosarcina acetivorans*. Acetate-dependent growth of a Δ*cam* mutant showed an extended lag phase, lower final cell density, and metabolized acetate to a threshold of 20.0 mM compared to 1.0 mM for wild-type. Molar growth yields (*Y*_methane_) were substantially greater for wild-type compared to the mutant. In contrast, growth parameters were identical for the methanol-grown wild-type and mutant. Rates of methane formation in resting cell suspensions containing 20.0 mM acetate were significantly less in the mutant *versus* wild-type and dependent on the presence of CO_2_. Rates for the wild-type decreased with increasing pH that was more pronounced for the mutant. CA activity was 100-fold greater in the membrane *versus* soluble fraction of acetate-grown cells. Addition of a surrogate CA stimulated acetate-dependent methanogenesis in resting cell suspensions of the mutant. The results support a role for Cam to supply protons for symport of acetate by the AceP symporter that also optimizes and facilitates growth at low acetate concentrations and high pH values encountered in the marine environment where *M. acetivorans* was isolated.

**Significance Statement:** Although CA plays major physiological roles in the domains *Eukarya* and *Bacteria*, a role has not been reported for the domain *Archaea* in which methanogens comprise the major group with abundant genomic annotations for CAs. Acetotrophic methanogens account for most of the methane produced in Earth’s biosphere where it is a major greenhouse gas. Although the biochemistry of the conversion of acetate to methane and carbon dioxide is well known, little is understood of acetate transport. The finding that CA has multiple roles facilitating thermodynamically constrained growth of a model marine acetotrophic methanogen has implications for advancing ecological understanding of the methane cycle that impacts global warming and climate change. Finally, the work is an introduction to anticipated physiological roles of CAs in the domain *Archaea* for which genomic annotations are abundant.

## Introduction

Carbonic anhydrase (CA) catalyzes the reversible hydration of CO_2_ to bicarbonate and a proton (Eq. 1), and is among the most widely distributed enzymes in all domains of life for which seven

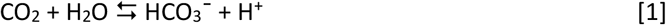

distinct subclasses are identified (α, β, γ, δ, ζ, η, θ and ι-CAs) (1, 2). CAs have been extensively investigated from the domain *Eukarya* where they serve multiple functions in terrestrial plants and mammals (3). CA is also essential for CO_2_ acquisition and growth of diverse marine phototrophs from the domains *Bacteria* and *Eukarya*. Marine diatoms, for example, contribute an estimated 20% of the global primary productivity dependent on CA for CO_2_ acquisition and concentration at the low levels present in oceans (4-6).

Although equally ubiquitous in the domains *Bacteria* and *Archaea*, and many biochemically characterized, few functions have been assigned apart from CO_2_ acquisition and concentration in phototrophs from the domain *Bacteria* (2, 7-9). Methane-producing species (methanogens) comprise a major group within the domain *Archaea* for which genomic annotations of CAs are abundant (10-14). CA activity in the acetate-grown methanogen *Methanosarcina barkeri* was reported in 1989 and remains the only documentation of CA in the domain *Archaea* (15). Remarkably, a physiological function has not been identified for any CA from the domain *Archaea*.

Acetate-utilizing (acetotrophic) methanogens (Eq. 2) of the domain *Archaea* are participants in

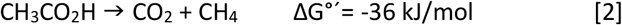

syntrophic microbial partnerships that decompose complex biomass to methane, an essential component of the global carbon cycle (16). The process accounts for approximately two-thirds of the estimated 400-500 Tg of biogenic methane produced yearly (17, 18). Most is oxidized to CO_2_ by aerobic and anaerobic methanotrophs. The remaining methane escapes to the upper atmosphere where the global warming potential is 30-fold greater than CO_2_; thus, acetotrophic methanogens account for a significant share of global warming and climate change (19). Indeed, it is proposed that evolution of marine acetotrophic methanogens produced a methanogenic burst in the end-Permian carbon cycle effecting global warming that triggered Earth’s greatest mass extinction (20).

Although major players in Earth’s methane cycle, the only known genera of acetotrophic methanogens are *Methanosarcina* and *Methanosaeta* for which the physiology has been extensively investigated except roles for CA. Additionally, exceptionally few studies of acetate transport in methanogens are reported (21, 22). *M. acetivorans* is the model marine acetotrophic methanogen for which the gamma class CA (Cam) is hypothesized to supply protons to AceP (locus MA_4008) shown to transport acetate by an acetate/proton symport mechanism (21). Furthermore, AceP displays 80% identity to the AceP homolog (locus MM_0903) of *Methanosarcina mazei* for which analyses of wild-type and the deletion mutant show it essential for acetotrophic growth (22). Finally, AceP is up regulated 125-fold when *M. acetivorans* is grown with acetate *versus* methanol (23). These reports indicate that AceP from *M. acetivorans* is responsible for proton-dependent symport of acetate during acetotrophic growth.

Herein is described an investigation of wild-type and a Δ*cam* mutant of *M. acetivorans* supporting multiple roles for Cam facilitating thermodynamically constrained acetotrophic growth with implications for survival and proliferation in the native marine environment.

## Results

Cam from *M. acetivorans* shares 87% identity (Figure S1) with the founding member of the gamma class CA from *M. thermophila* which is biochemically characterized although a physiological function has not been assigned (12, 24-29). Sequence comparisons show strict conservation of catalytically and structurally relevant residues (Figure S1).

Growth was evaluated for wild-type and a Δ*cam* mutant cultured with either acetate or methanol in HEPES-buffered media and a N_2_ atmosphere absent of CO_2_ to examine the role of Cam in the transport of acetate in *M. acetivorans* (Fig. 1). Acetate-grown wild-type achieved a final cell density greater than the Δ*cam* mutant accompanied by a lag phase approximately two days shorter than the mutant (Figure 1A,1B). The wild-type produced a proportional 15-20% more methane with a greater molar growth yield (*Y*_methane_) during exponential growth (2 ± 0.1 *versus* 2.6 ± 0.3 g dry weight/mol methane, respectively). Furthermore, the wild-type metabolized acetate to a threshold level of 1.96 ± 0.75 mM compared to 18.8 ± 2.8 mM for the Δ*cam* mutant (Figure 1C) consistent with the relative amounts of methane produced (Figure 1B). These results show that Cam is essential for optimum growth with acetate. In contrast, both wild-type and the Δ*cam* mutant grew equally well with methanol (Figure 2) in terms of cell density, methane production, and *Y*_methane_ (5 ± 0.3 and 5 ± 0.4, respectively) showing no significant role for Cam in methylotrophic growth and methanogenesis. The differences in acetotrophic growth were more striking when wild-type and the Δ*cam* mutant were grown with 50.0 *versus* 100.0 mM acetate indicating a role for Cam to facilitate growth at low concentrations of acetate (Figure 3).

**Figure 1.**
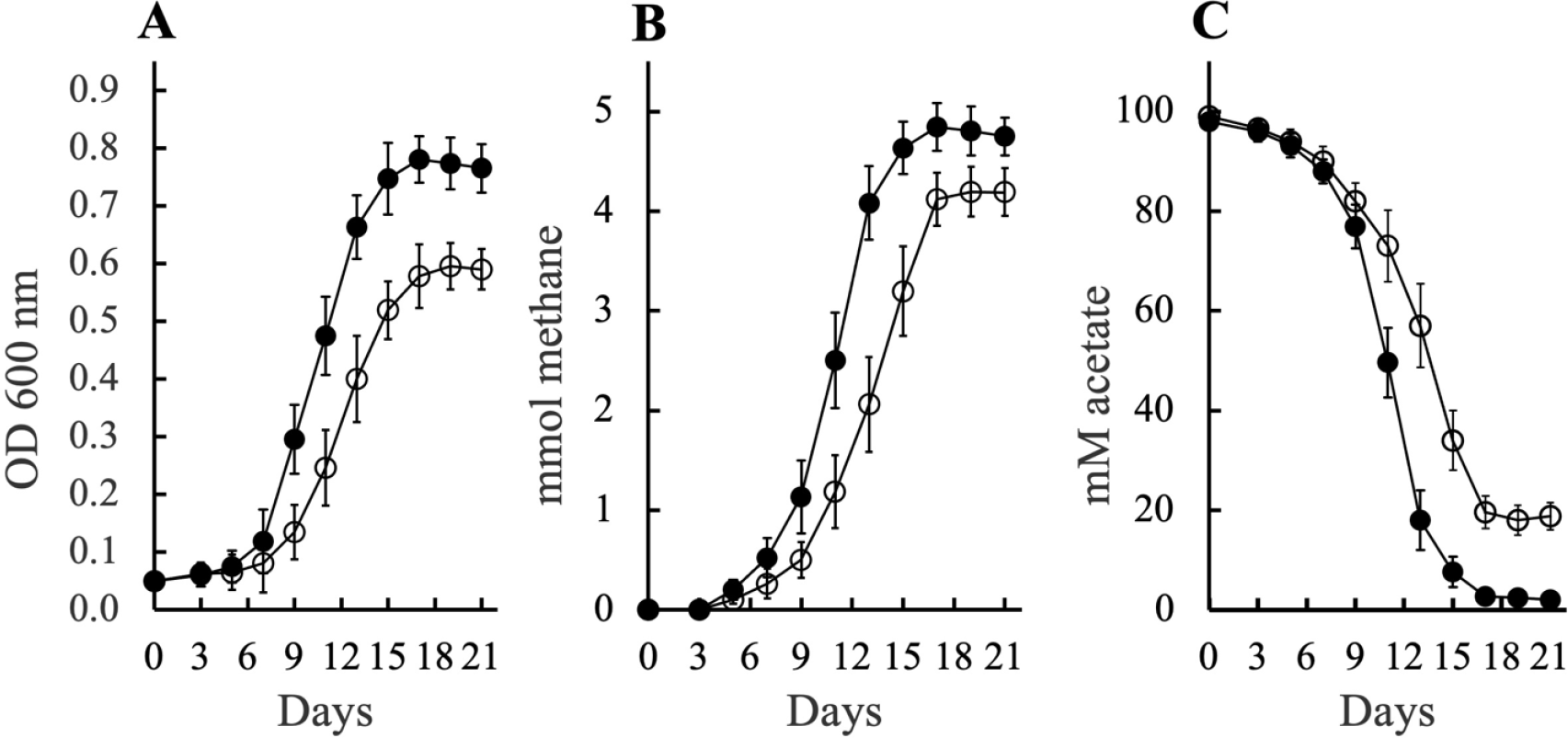
Growth of *M. acetivorans* with 100.0 mM acetate at pH 6.8. Wild-type (●), Δ*cam* mutant **(○)**. Bars show the mean and standard deviation of eight biological replicates.

**Figure 2.**
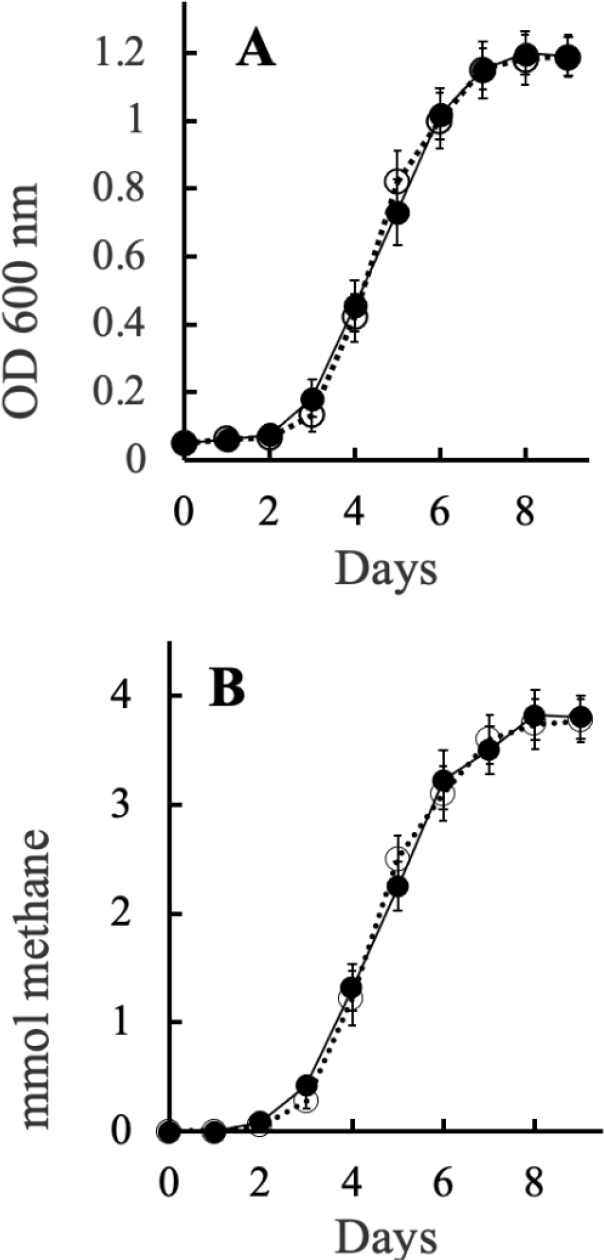
Growth of *M. acetivorans* with 100.0 mM methanol. A, growth. B, methane production. Wild-type (●), Δ*cam* mutant (○). Bars show the mean and standard deviation of eight biological replicates.

**Figure 3.**
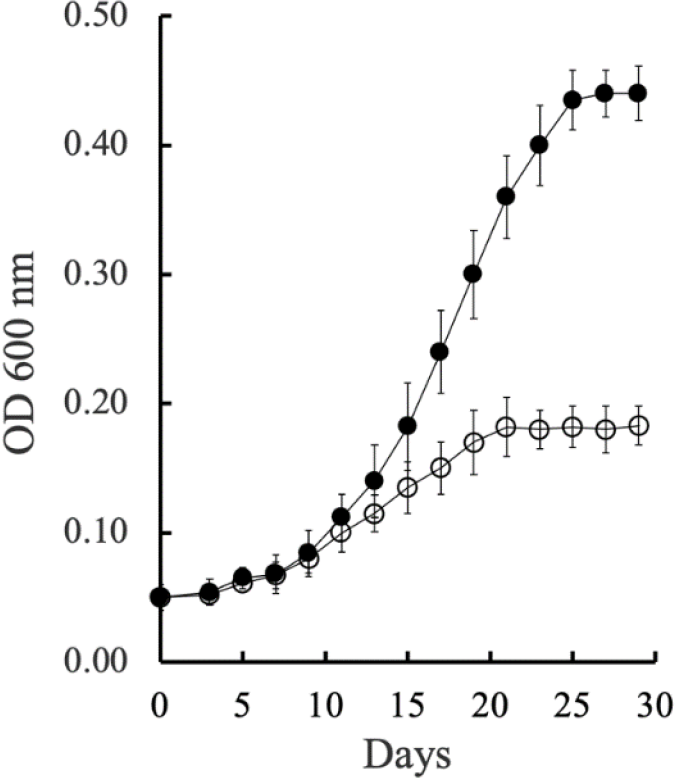
Growth of *M. acetivorans* with 50.0 mM acetate. Wild-type (●), Δ*cam* mutant (○). Bars show the mean and standard deviation of eight biological replicates.

The results shown In Table 1 support a proton symport mechanism for acetate transport. Acetate-dependent methanogenesis in resting cell suspensions was severely inhibited by addition of the protonophore carbonyl cyanide-m-chlorophenyl-hydrazone (CCCP) whereas no significant inhibition was observed with addition of the sodium ionophore ETH157. The addition of ETH157 showed the opposite effect for methanol-grown cells that require a sodium gradient to drive methyl transfer in the oxidative branch of the pathway for methanogenesis from methanol (30). Biochemical analyses show AceP from *M. acetivorans* transports acetate by a proton symport mechanism, mutant studies show an AceP homolog is required for acetate transport in *M. mazei*, and AceP is up regulated 125-fold in acetate-*versus* methanol-grown *M. acetivorans*. These results, and the results shown in Table 1, establish a role for AceP in the proton symport of acetate.

**Table 1.**
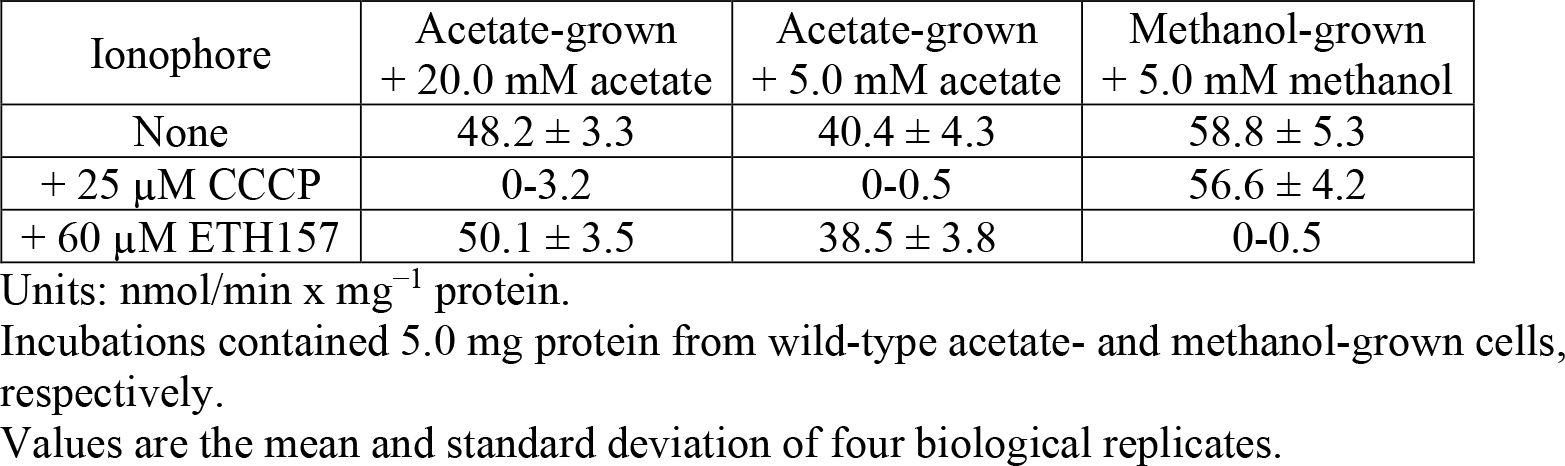
Effect of proton and sodium ionophores on acetate-dependent methanogenesis in resting cell suspensions of *M. acetivorans*.

The hypothesized role of Cam to supply protons to AceP was investigated in short term experiments with resting cell suspensions in the presence or absence of CO_2_ required for Cam activity (Eq. 2). Figure 4 shows declining rates of methanogenesis in response to decreasing concentrations of acetate for both wild-type and the Δ*cam* mutant in the presence of 0.15 atm CO_2_. The decline was substantially greater for the Δ*cam* mutant, and no significant methane was detected with 5.0 mM acetate for the mutant. Notably, no methane was detected for wild-type or the Δ*cam* mutant at all concentrations of acetate in the absence of added CO_2_ (Table S1). This result, combined with the results shown in Figure 1C, show a more acute role for Cam at lower concentrations of acetate and a CO_2_ requirement for CA activity (Eq. 2) consistent with the proposed role for Cam to supply protons for acetate symport by AceP. Methane production from methanol was not statistically different for wild-type and the Δ*cam* mutant with or without 0.15 atm CO_2_ (Table S1) which supports roles for Cam specific to acetotrophic methanogenesis and growth.

**Figure 4.**
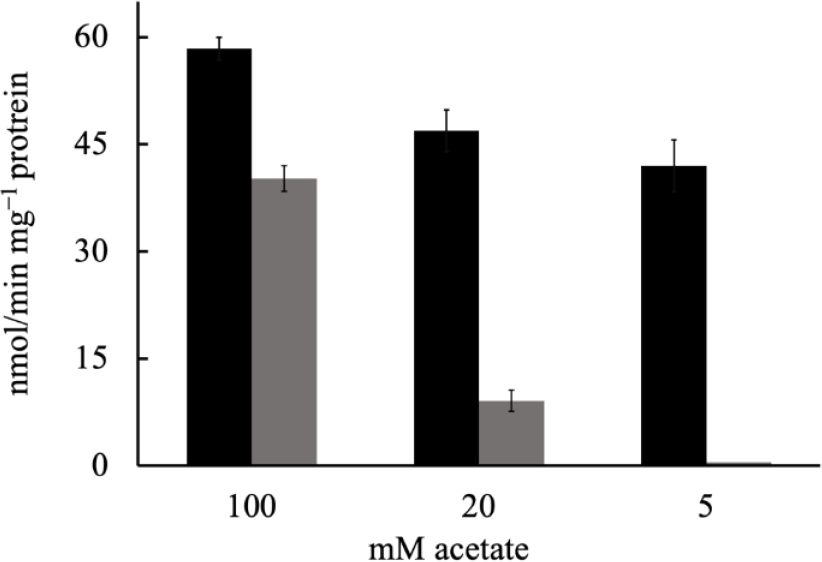
Rates of methanogenesis from resting cell suspensions of *M. acetivorans* dependent on the concentration of acetate. Suspensions (pH 6.8) contained 0.15 atm CO_2_. Wild-type (black), *Δcam* mutant strain (gray). No methane was detected in controls minus substrate. Bars show the mean and standard deviation of four biological replicates. The units are: nmol of methane produced x min^-1^ x mg^-1^ of total protein.

Further support for the role of Cam to supply protons to AceP for symport of acetate is shown in Table 2. The rate of methane production in resting cell suspensions containing 100.0 mM acetate was greater for wild-type compared to the mutant at all pH values tested. These rates decreased with increasing pH consistent with an acetate/symport mechanism. Although methanogenesis for the Δ*cam* mutant also decreased with increasing pH, the rates were substantially lower with no detectible methane produced at pH 7.4 and 7.6. The results are consistent with the hypothesized role of Cam to replace the supply of protons to AceP that diminish with increasing pH. However, Figure 5 shows equal growth of wild-type and the Δ*cam* mutant with 100.0 mM acetate at pH 6.0 in contrast to growth with 100.0 mM acetate at pH 6.8 (Figure 1) for which the wild-type dominated. The result is consistent with a greater proportion of acetic acid at pH 6.0 which can freely diffuse across the membrane thereby minimizing the requirement for CA activity to supply protons for symport by AceP. However, the result in Figure 5 conflicts with the result in Table 2 where the rate of methanogenesis is less for the mutant compared to wild-type at pH 6.0. This apparent anomaly is possibly the result of resting cells uncoupled from growth for which the rate of methanogenesis is greater than transport requiring Cam.

**Table 2.** Effect of pH on the rate of methanogenesis from acetate in resting cell suspensions of *M. acetivorans* wild-type and the *Δcam* mutant. Suspensions contained 100.0 mM acetate and 5.0 mg cell protein from acetate-grown wild-type or the *Δcam* mutant.

| pH | wild-type | *Δcam* mutant |
| --- | --- | --- |
| 6.0 | 139 ± 10 | 92 ± 8 |
| 6.8 | 58.4 ± 1.6 | 40.2 ± 1.8 |
| 7.4 | 30 ± 3.5 | < 4 |
| 7.6 | 16 ± 2.5 | < 4 |

Units: nmol/minute x mg<sup>-1</sup> protein.
Values are the mean and standard deviation of four biological replicates.

**Figure 5.**
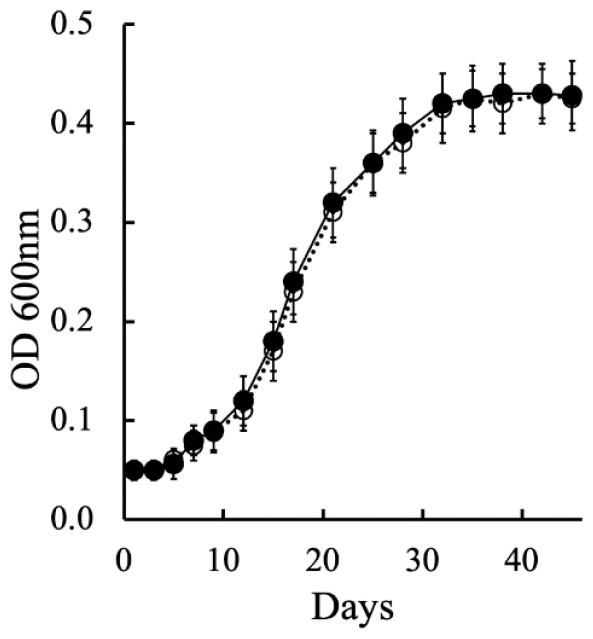
Growth of *M. acetivorans* at pH 6.0 and with 100.0 mM acetate. Wild-type (●) Δ*cam* mutant (○). Bars show the mean and standard deviation of five biological replicates.

The hypothesized role for Cam requires at least a portion located on the outer aspect of the cytoplasmic membrane to supply protons to AceP for symport of acetate. Figure 6 shows the rate of methanogenesis from acetate in resting cell suspensions of the Δ*cam* mutant enhanced 65% by addition of commercially available α-CA suggesting a role for CA activity outside the cytoplasmic membrane for efficient transport of acetate. Support for a membrane-associated location of Cam is shown in Table 3. CA activity in the membrane fraction of acetate-grown cells was 100-fold greater than the cytosolic fraction. Acetate kinase (Ack) is a cytosolic enzyme for which enzymatic activity was absent in the membrane fractions that confirmed no significant contamination with cytosolic enzymes (31). CA activity was not detected in the cell-free extract of Δ*cam* mutant cells grown with acetate or methanol showing Cam is responsible for CA activity in the membrane. Furthermore, CA activity in the membrane fraction from methanol-grown cells is 36-fold less when *cam* is down regulated relative to acetate-grown cells (32). Together, these results indicate Cam is membrane-bound.

**Table 3.** Carbonic anhydrase activity in the membrane and cytosolic fractions of wild-type *M. acetivorans*.

| Enzyme activity | Acetate-grown |  | Methanol-grown |  |
| --- | --- | --- | --- | --- |
|  | Membrane fraction | Cytosolic fraction | Membrane fraction | Cytosolic fraction |
| <sup>a</sup>Carbonic anhydrase | 12.93 ± 0.05 | 0.12 ± 0.01 | 0.36 ± 0.01 | ND |
| <sup>b</sup>Acetate kinase | ND | 0.72 ± 0.02 | ND | 0.14 ± 0.06 |

<sup>a</sup>Units are µmol H<sup>+</sup> produced/minute x mg protein<sup>-1</sup>.
<sup>b</sup>Units are µmol NADH consumed/minute x mg protein<sup>-1</sup>.
ND: Not detected. Detection limit: 0.01 µmol H<sup>+</sup> produced or 0.05 nmol NADH consumed.
Values are the mean and standard deviation of three biological replicates.

**Figure 6.**
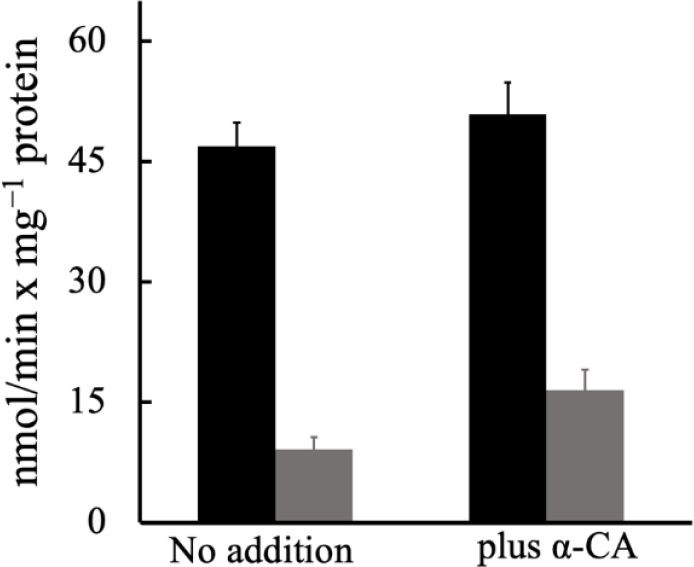
Rates of methane production from resting cell suspensions of *M. acetivorans* in the presence or absence of an alpha class carbonic anhydrase. Suspensions (pH 6.8) were incubated at 37ºC and contained 20.0 mM acetate and 0.15 atm of CO_2_. Wild-type (black), *Δcam* mutant (grey). Bars show the mean and standard deviation of four biological replicates. The units are: nmol of methane produced x min^-1^ x mg^-1^ of total protein.

## Discussion

### Physiological roles

The findings establish roles for Cam in acetotrophic growth that govern the lag phase, growth yield, pH dependence, and threshold concentration of acetate. The results shown in Figure 1 are consistent with the proposed role of Cam to supply protons to AceP. Both wild-type and the Δ*cam* mutant grew with 100 mM acetate at pH 6.8 in the absence of CO_2_ essential for Cam activity indicating that the proportion of acetic acid is sufficient for diffusion across the cytoplasmic membrane to initiate growth. However, as growth progresses and product CO_2_ accumulates (Eq. 2), the Cam activity of wild-type contributes protons to AceP which shortens the lag phase relative to the Δ*cam* mutant. As growth continues, the proportion of acetic acid becomes limiting and acetate symport by AceP is increasingly dependent on Cam for optimal growth and lowering the threshold concentration of acetate uptake. The pH dependent growth of wild-type compared to the mutant further supports the proposed role for Cam to supply protons to AceP. At low pH, the proportion of acetic acid is sufficient for diffusion across the membrane to support growth. However, at increasing pH values, the proportion of acetic acid becomes limiting with greater reliance on Cam to supply protons to AceP for symport. Enhancement of growth yield (*Y*_methane_) dependent on Cam is attributed to the contribution of protons for acetate symport otherwise supplied by the proton gradient driving ATP synthesis and growth. Finally, the sodium ionophore ETH157 had no effect on methanogenesis from acetate in resting cell suspensions which rules out a role for sodium and the Mrp sodium/proton exchanger to supply protons to AceP (33). These proposed roles for Cam predict that Cam and AceP form a complex in which at least a portion of Cam is located on the outer aspect of the cytoplasmic membrane (Figure 7). A location for Cam at the outer aspect of the cytoplasmic membrane is supported by the finding that CA activity is confined to Cam and the membrane of *M. acetivorans*, and that a surrogate CA enhanced the rate of methanogenesis when added to resting cell suspensions of the Δ*cam* mutant. The cellular location is further supported by an N-terminal sequence with 74% similarity and 47% identity (Figure S1) to the post translational cleaved leader sequence shown for the Cam homolog from *M. thermophila* (14). Finally, a signal peptide (resides 1-26) for *M. acetivorans* Cam was confirmed and a continuous non-cytoplasmic sequence (residues 27-247) was predicted (Figure S2) with high probability using the Phobius method of sequence analysis (https://phobius.sbc.su.se/) consistent with a membrane location (34).

**Figure 7.**
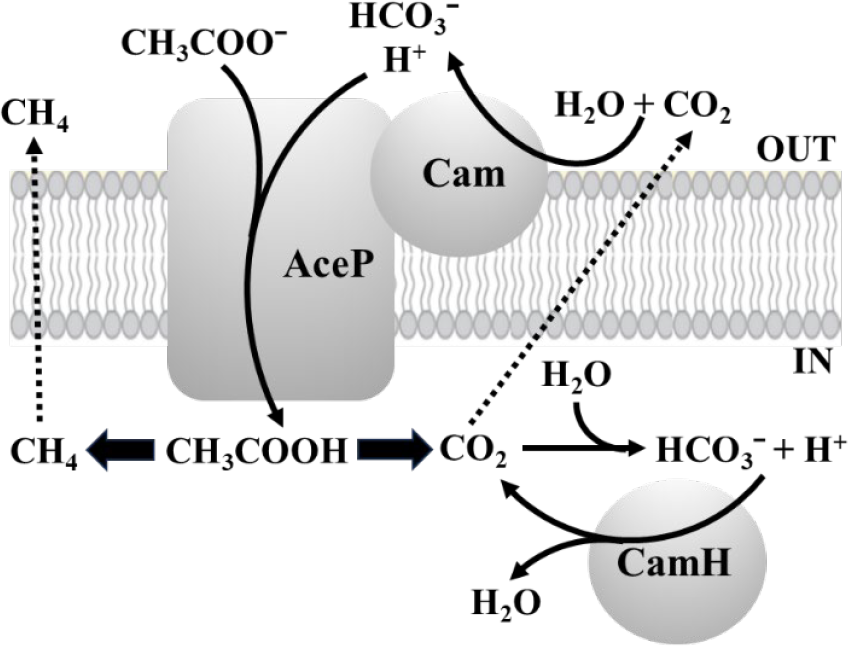
Proposed roles for Cam, AceP, and CamH enabling acetate transport and growth in *M. acetivorans*. Dashed arrows indicate diffusion and thick arrows metabolism.

The proposed spatial juxtaposition of Cam and AceP would assure efficient transfer of protons to AceP favoring the direction of CO_2_ hydration by reversible CA activity (Eq. 1). Annotations for genes encoding bicarbonate transporters are absent in the *M. acetivorans* genome, consistent with diffusion of CO_2_ from the cytoplasm to outside the membrane (Figure 7).

The genome of *M. acetivorans* is annotated with a gene (MA1098) 85% identical to *camH* of *M. thermophila* (Fig. S4) which is the founding member of a subclass of the gamma class that has been biochemically characterized (11). It is proposed that CamH of *M. thermophila* is cytosolic and involved in growth with acetate, but not with methanol (11, 35). The *camH* gene of *M. acetivorans* is up-regulated 20-fold in cells grown with acetate *versus* methanol suggesting CamH has a role during acetotrophic growth (35). Like *M. thermophila*, the deduced sequence of CamH from *M. acetivorans* is missing the leader sequence of Cam consistent with a cytosolic location (Figure S4). Although no CA activity was detected in the cell-free extract of the mutant, the result does not necessarily preclude an active role. The products of acetate metabolism are methane and CO_2_ for which the latter is hydrated to bicarbonate in a spontaneous reversable reaction (Eq. 1). A potential role for CamH is to shift the reaction towards CO_2_ for diffusion across the membrane where it interacts with Cam to supply protons to AceP (Figure 7). Indeed, comparison of the homologous enzymes from *M. thermophila* reveal the *K*_*m*_ (CO_2_) and catalytic efficiency (*K*_*cat*_/*K*_*m*_) for CamH 10- and 4-fold less than Cam (11).

### Ecological implications

Dissolved inorganic carbon in sea water (Eq. 3) is governed by atmospheric CO_2_ at 0.04% (36). However, *M. acetivorans* is routinely grown in high-salt media (HS) with a bicarbonate buffer system comprised of 45 mM Na_2_CO_3_ and an atmosphere of 20% CO_2_ balanced with 80% N_2_ (37). Thus, growth of *M. acetivorans* with the HEPES buffer system

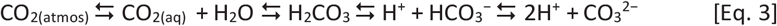

facilitated an understanding of roles CO_2_ and CA play in acetotrophic growth of *M. acetivorans*. The proposed multiple roles Cam and CamH play in acetotrophic growth potentially provide *M. acetivorans* the means for survival and proliferation in the native marine environment by lowering the concentration of acetate accessible for growth and enhancing growth by contributing protons for acetate symport otherwise supplied by the proton gradient driving ATP synthesis. Removal of CO_2_ by conversion to membrane impermeable bicarbonate is yet another mechanism proposed to improve the thermodynamics for growth (38). These roles are important as the available energy for growth is marginal under standard conditions of equimolar reactants and products (Eq. 2) which are potentially more favorable for growth than that encountered in the native environment. The proposed roles for Cam should be accentuated in the marine environment where the mean pH of seawater is 8.0 and acetic acid is less abundant (36).

### Conclusions

The work complements previous investigations of a model marine methanogen to provide a comprehensive understanding of thermodynamically constrained growth in the environment with implications for greater ecological understanding of the marine methane cycle. The work also expands the multiple roles carbonic anhydrase plays in the diverse life and chemistry of Earth’s oceans. Finally, the work is the first to recognize a physiological role for any carbonic anhydrase in the domain *Archaea* for which genomic annotations are replete.

## Materials and Methods

### Materials

Pyruvate kinase, L-lactate dehydrogenase and phosphoenolpyruvate were purchased from Roche (Manheim, Germany). HEPES, Tris-HCl, sodium acetate, ATP, NADH, acetate kinase from *M. thermophila* and alpha carbonic anhydrase from bovine erythrocytes were purchased from Sigma-Aldrich (St Louis, MO, USA). Absolute methanol was of analytical grade purchased from Thermo Fisher Scientific Inc. (Pittsburgh, PA, USA).

### Growth

Growth of *M. acetivorans* strain C2A wild-type and the Δ*cam* mutant strain was in high-salt media (37) modified by replacing the sodium bicarbonate/CO_2_ buffer with 45 mM HEPES (4-(2-hydroxyethyl)-1-piperazineethanesulfonic acid), and an atmosphere of 100% N_2_. The pH was adjusted to 6.8 with dilute HCl.

Trimethylamine-grown wild-type and mutant strains were adapted to growth with 100.0 mM methanol by 10 serial transfers into fresh media. Methanol-grown wild-type was adapted to growth with acetate by serial dilution of stationary phase cultures (1:100) into fresh medium containing 75.0 mM methanol and 25.0 mM acetate. With each subsequent passage, the concentration of acetate was increased, and the concentration of methanol was decreased until growth with 100.0 mM acetate was achieved. The wild-type strain was considered adapted after 10 passages in media containing 100.0 mM acetate. The Δ*cam* mutant strain required a more stringent adaptation protocol. The Δ*cam* strain previously grown on methanol was adapted to acetate by serially diluting stationary phase cells into fresh media containing 10.0 mM acetate and 90.0 mM methanol. On each subsequent passage, the concentration of acetate was increased by 10.0 mM and the concentration of methanol was decreased by 10.0 mM until growth with 100.0 mM acetate was achieved. The Δ*cam* strain was considered adapted to acetate after 10 passages into media containing 100.0 mM acetate.

### Construction of the *cam* deletion (Δ*cam*) strain

The gene encoding *cam* (MA2536) from *M. acetivorans*, grown with 10.0 mM trimethylamine and 50.0 mM acetate, was amplified 434bp upstream and 510bp downstream of the structural gene with primers 234 (5’-CCCTAGTGTCTCGTCAATTCTTAAA-3’) and 235 (5’-CCTACAAATTAGTCTGCTTGAACGA-3’) by polymerase chain reaction (PCR) as described previously (39). A reaction was set up with an AmpliTaq kit (Invitrogen) using 1× PCR buffer II, 1.5 mM MgCl_2_, 1.6 mM deoxynucleotide triphosphate mix, 1 μg *M. acetivorans* C2A genomic DNA, 2 pmol of each primer, 2.5 U AmpliTaq DNA polymerase in a 50 μl total volume. Conditions for PCR were as follows: initial denaturation at 94°C for 5 min, followed by 30 amplification cycles of 94°C for 30s, 55°C for 1 min, and 72°C for 3 min. The PCR product was ligated into PCR 4.0 cloning vector (Invitrogen TOPO cloning kit) to generate pEA156. Plasmid pEA156 was digested with restriction enzymes *Mfe*I and *BstB*I resulting in a deletion of 442bp from the *cam* gene. A serCP::proC cassette PCR amplified from pJK88 (40) with primers 251 (5’-CCACCTTTTTTCCT<u>CAATTG</u>GTGAACTC-3’) (*Mfe*I site underlined) and 252 (5’-CGTAACCGC<u>TTCGAA</u>GTGGCGGCC-3’) (*BstB*I site underlined), as described above, and ligated into the unique *Mfe*I and *BstB*I sites in *cam*, which generated pEA162. The *cam*::serCP::proC deletion insert was amplified by PCR with primers 234 and 235, and 4 μg of the PCR product was used to transform competent *M. acetivorans* strain WWM24, a proline auxotroph, as described previously (41). The transformed cultures were selected on solidified medium without addition of proline and screened for insertion of the *cam* deletion with primers 268 (5’-GCCCTTCTGGGTTCAGAACCC-3’) AND 269 (5’-CACCGGGAGGCACCACCATTGAAGC-3’), which amplified 909 bp upstream of *cam* structural gene and the 3’ end of *pro*C. The correct construct was confirmed in three clones by sequencing and one clone (KSC62) was selected for experiments. Antibody probing of the mutant showed no cross reactivity with anti-Cam antisera (35).

### Analytical

Methane was determined by gas chromatography as previously reported (42). Acetate in spent media was determined spectrophotometrically by conversion of ATP and acetate to ADP and acetyl phosphate coupled to oxidation of NADH to NAD^+^ by pyruvate kinase and lactate dehydrogenase (43). Cultures were centrifuged at 3,500 x *g* for 10 minutes and the cell-free supernatant solution diluted with 50.0 mM HEPES buffer (pH 7.5) containing 10.0 mM MgCl_2_ and 1.0 mM EGTA (ethylene glycol tetra acetic acid). The assay reaction mixture contained 5.0 mM ATP, 2.0 mM phosphoenolpyruvate, 0.25 mM NADH. Acetate kinase, pyruvate kinase and lactate dehydrogenase were added in excess (0.5 units of each) to ensure the complete phosphorylation of acetate coupled to NADH oxidation.

Molar growth yields (*Y*_methane_= g dry weight/mol) were determined at days 8 and 19 for cells grown with acetate and methanol, respectively. Dry weight was estimated using the previously published value of one OD_600nm_ unit corresponding to 0.32 g/liter (44).

Protein was determined with a modified Biuret method (45). Briefly, 1 ml culture samples were incubated with 10% (w/v) trichloroacetic acid at 4°C for 2 h and centrifuged at 14,462 x *g* for 5 minutes. The pellets were resuspended in 1 ml deionized water. Protein in the resuspended pellet was determined using bovine serum albumin as the standard.

### Enzyme assays

The spectrophotometric assay for acetate kinase activity was as previously described (46). Carbonic anhydrase activity was measured electrometrically in the direction of CO_2_ hydration as previously described except performed anaerobically in stoppered serum vials (10 ml) containing 5 ml of 100.0 mM Tris buffer (pH 8.2) and an atm of N_2_ (14). Stoppers were adapted to the size of the pH electrode to maintain the anaerobic environment during measurements. The reaction was started by addition of protein samples. Activity of the alpha class carbonic anhydrase control was 166.2 μmol H^+^ produced/minute x mg protein^-1^.

Cell fractions. Membrane and cytosolic fractions were obtained from cell-free extract using a discontinuous sucrose gradient as described (47). The membrane fraction was collected from the 30/70 % sucrose interface and the cytosolic fraction from the membrane-depleted supernatant and stored at minus 80°C until use.

### Resting cell suspensions

Wild-type and the Δ*cam* mutant grown with 100.0 mM acetate or 100.0 mM methanol were harvested in the late exponential phase of cell growth by centrifugation of the 250 ml suspension as described above. Cells were washed with 200 ml of TME buffer (50.0 mM Tris, 20.0 mM MgCl_2_, 1.0 mM EGTA) at pH 7.5 to remove remnant energy sources. The protein was quantified, and the cell pellet re-suspended in 5 ml of oxygen-free HEPES-buffered media (pH 6.8) contained in stoppered serum bottles (10 ml). Cell suspensions (5.0 mg protein) were pre-incubated for 5 min at 37_o_C without agitation. The headspace (1.0 Atm.) was 85% N_2_ and 15% CO_2_, or 100% N_2_. After pre-incubation, substrate was added at the indicated concentration and incubated at 37°C. The increase of methane determined at 15, 30, 45, and 60 min was linear.

## Acknowledgments

This work was supported by the Stanley R. Person endowment of Penn State to J.G.F. and the Division of Chemical Sciences, Geosciences, and Biosciences, Office of Basic Energy Sciences of the U.S. Department of Energy through grants DE-SC0018035 to J.G.F and DE-FG02-93ER20106 to K.R.S.

## References

1. D. Hewett-Emmett, “Evolution and distribution of the carbonic anhydrase gene families” in The Carbonic Anhydrases, W. R. Chegwidden, N. D. Carter, Y. H. Edwards, Eds. (Birkhauser Verlag, Basel, 2000), pp. 29–76.

2. B. C. Tripp, K. S. Smith, J. G. Ferry, Carbonic anhydrase: new insights for an ancient enzyme. J Biol Chem 5, 5 (2001).

3. R. P. Henry, Multiple roles of carbonic anhydrase in cellular transport and metabolism. Annu. Rev. Physiol. 58, 523–538 (1996).

4. E. V. Armbrust, The life of diatoms in the world’s oceans. Nature 459, 185–192 (2009).

5. P. G. Falkowski et al., The evolution of modern eukaryotic phytoplankton. Science 305, 354–360 (2004).

6. J. J. Pierella Karlusich, C. Bowler, H. Biswas, Carbon dioxide concentration mechanisms in natural populations of marine diatoms: insights from tara oceans. Front. Plant Sci. 12, 657821 (2021).

7. C. Capasso, C. T. Supuran, An overview of the alpha-, beta- and gamma-carbonic anhydrases from Bacteria: can bacterial carbonic anhydrases shed new light on evolution of bacteria? J Enzyme Inhib Med Chem 30, 325–332 (2015).

8. C. T. Supuran, C. Capasso, An Overview of the Bacterial Carbonic Anhydrases. Metabolites 7 (2017).

9. R. S. S. Kumar, J. G. Ferry, “Prokaryotic Carbonic Anhydrases of Earth’s Environment” in Carbonic Anhydrase: Mechanism, Regulation, Links to Disease, and Industrial Applications, S. C. Frost, R. McKenna, Eds. (Springer Science Business Media, Dordrecht, 2014), vol. Subcellular Biochemistry 75, chap. 5, pp. 77–87.

10. J. G. Ferry, “Zinc and Iron, Gamma and Beta Class, Carbonic Anhydrases of the Domain Archaea” in Encyclopedia of Metalloproteins, V. N. Uversky, R. H. Kretsinger, E. A. Permyakov, Eds. (Springer-Verlag, Berlin Heidelberg, 2013), pp. 2380–2385.

11. S. A. Zimmerman, J. F. Tomb, J. G. Ferry, Characterization of CamH from Methanosarcina thermophila, founding member of a subclass of the γ class of carbonic anhydrases. J. Bacteriol. 192, 1353–1360 (2010).

12. B. C. Tripp, C. B. Bell, F. Cruz, C. Krebs, J. G. Ferry, A role for iron in an ancient carbonic anhydrase. J. Biol. Chem. 279, 6683–6687 (2004).

13. K. S. Smith, J. G. Ferry, A plant-type (beta-class) carbonic anhydrase in the thermophilic methanoarchaeon Methanobacterium thermoautotrophicum. J. Bacteriol. 181, 6247–6253 (1999).

14. B. E. Alber, J. G. Ferry, A carbonic anhydrase from the archaeon Methanosarcina thermophila. Proc. Natl. Acad. Sci. USA 91, 6909–6913 (1994).

15. M. Karrasch, M. Bott, R. K. Thauer, Carbonic anhydrase activity in acetate grown Methanosarcina barkeri. Arch. Microbiol. 151, 137–142 (1989).

16. J. G. Ferry, “Acetate: A Key Intermediate during the Anaerobic Degradation of Organic Matter” in Acetate: Versatile Building Block of Biology and Chemistry, D. A. Sanders, Ed. (Nova Science, Hauppauge, 2013), chap. 1, pp. 1–14.

17. R. Conrad, The global methane cycle: recent advances in understanding the microbial processes involved. Environ. Microbiol. Rep. 1, 285–292 (2009).

18. R. K. Thauer, A. K. Kaster, H. Seedorf, W. Buckel, R. Hedderich, Methanogenic archaea: ecologically relevant differences in energy conservation. Nat. Rev. Microbiol. 6, 579–591. doi: 10.1038/nrmicro1931 (2008).

19. R. G. Derwent, Global warming potential (GWP) for methane: monte carlo analysis of the uncertainties in global tropospheric model predictions. Atmosphere 11, 486 (2020).

20. D. H. Rothman et al., Methanogenic burst in the end-Permian carbon cycle. Proc. Natl. Acad. Sci. U. S. A. 111, 5462–5467 (2014).

21. D. Ribas et al., The acetate uptake transporter family motif “NPAPLGL(M/S)” is essential for substrate uptake. Fungal. Genet. Biol. 122, 1–10 (2018).

22. C. Welte, L. Kroninger, U. Deppenmeier, Experimental evidence of an acetate transporter protein and characterization of acetate activation in aceticlastic methanogenesis of Methanosarcina mazei. FEMS Microbiol. Lett. 359, 147–153 (2014).

23. L. Rohlin, R. P. Gunsalus, Carbon-dependent control of electron transfer and central carbon pathway genes for methane biosynthesis in the Archaean, Methanosarcina acetivorans strain C2A. BMC Microbiol. 10, 62 (2010).

24. B. E. Alber et al., Kinetic and spectroscopic characterization of the gamma-carbonic anhydrase from the methanoarchaeon Methanosarcina thermophila. Biochemistry 38, 13119–13128 (1999).

25. B. C. Tripp, J. G. Ferry, A structure-function study of a proton transport pathway in a novel γ-class carbonic anhydrase from Methanosarcina thermophila. Biochemistry 39, 9232–9240 (2000).

26. B. C. Tripp, C. Tu, J. G. Ferry, Role of arginine 59 in the γ-class carbonic anhydrases. Biochemistry 41, 669–678. (2002).

27. S. A. Zimmerman, J. G. Ferry, Proposal for a hydrogen bond network in the active site of the prototypic γ-class carbonic anhydrase. Biochemistry 45, 5149–5157 (2006).

28. S. Zimmerman et al., Role of Trp19 and Tyr200 in catalysis by the gamma-class carbonic anhydrase from Methanosarcina thermophila. Arch. Biochem. Biophys. 529, 11–17 (2013).

29. T. M. Iverson, B. E. Alber, C. Kisker, J. G. Ferry, D. C. Rees, A closer look at the active site of γ-carbonic anhydrases: High resolution crystallographic studies of the carbonic anhydrase from Methanosarcina thermophila. Biochemistry 39, 9222–9231 (2000).

30. U. Deppenmeier, The membrane-bound electron transport system of Methanosarcina species. J. Bioenerg. Biomembr. 36, 55–64 (2004).

31. D. J. Aceti, J. G. Ferry, Purification and characterization of acetate kinase from acetate-grown Methanosarcina thermophila. J. Biol. Chem. 263, 15444–15448 (1988).

32. L. Li et al., Quantitative proteomic and microarray analysis of the archaeon Methanosarcina acetivorans grown with acetate versus methanol. J. Proteome. Res. 6, 759–771 (2007).

33. R. Jasso-Chavez, E. E. Apolinario, K. R. Sowers, J. G. Ferry, MrpA functions in energy conversion during acetate-dependent growth of Methanosarcina acetivorans. J. Bacteriol. 195, 3987–3994 (2013).

34. L. Käll, A. Krogh, E. L. Sonnhammer, A combined transmembrane topology and signal peptide prediction method. J. Mol. Biol. 338, 1027–1036 (2004).

35. S. A. Zimmerman (2007) Understanding the biochemistry and physiology of gamma carbonic anhydrases in Methanosarcina thermophila. in Biochemistry and Molecular Biology (Pennsylvania State University), pp 1–161.

36. S. C. Doney, V. J. Fabry, R. A. Feely, J. A. Kleypas, Ocean acidification: the other CO2 problem. Ann. Rev. Mar. Sci. 1, 169–192 (2009).

37. K. R. Sowers, J. E. Boone, R. P. Gunsalus, Disaggregation of Methanosarcina spp and growth as single cells at elevated osmolarity. Appl. Environ. Microbiol. 59, 3832–3839 (1993).

38. J. S. Liu, I. W. Marison, U. von Stockar, Microbial growth by a net heat up-take: A calorimetric and thermodynamic study on acetotrophic methanogenesis by Methanosarcina barkeri. Biotechnol. Bioeng. 75, 170–180 (2001).

39. R. Jasso-Chavez, C. Diaz-Perez, J. S. Rodriguez-Zavala, J. G. Ferry, Functional role of MrpA in the MrpABCDEFG Na+/H+ antiporter complex from the archaeon Methanosarcina acetivorans. J. Bacteriol. 199, e00662–00616 (2017).

40. J. K. Zhang, A. K. White, H. C. Kuettner, P. Boccazzi, W. W. Metcalf, Directed mutagenesis and plasmid-based complementation in the methanogenic archaeon Methanosarcina acetivorans C2A demonstrated by genetic analysis of proline biosynthesis. J. Bacteriol. 184, 1449–1454 (2002).

41. W. W. Metcalf, J. K. Zhang, E. Apolinario, K. R. Sowers, R. S. Wolfe, A genetic system for Archaea of the genus Methanosarcina: Liposome-mediated transformation and construction of shuttle vectors. Proc. Natl. Acad. Sci. USA 94, 2626–2631 (1997).

42. D. Prakash, S. S. Chauhan, J. G. Ferry, Life on the thermodynamic edge: Respiratory growth of an acetotrophic methanogen. Sci. Adv. 5, 1–6 (2019).

43. P. M. Clarke, M. A. Payton, An enzymatic assay for acetate in spent bacterial culture supernatants. Anal. Biochem. 130, 402–405 (1983).

44. K. R. Sowers, M. J. K. Nelson, J. G. Ferry, Growth of acetotrophic, methane-producing bacteria in a pH auxostat. Curr. Microbiol. 11, 227–230 (1984).

45. A. G. Gornall, C. J. Bardawill, M. M. David, Determination of serum proteins by means of the Biuret reaction. J. Biol. Chem. 177, 751–766 (1948).

46. P. Iyer, J. G. Ferry, “Acetate kinase from Methanosarcina thermophila, a key enzyme for methanogenesis” in Microbial Enzymes and Biotransformations, J. L. Barredo, Ed. (Humana Press Inc., Totowa, NJ, 2005), vol. 17, pp. 239–246.

47. M. Wang, J. F. Tomb, J. G. Ferry, Electron transport in acetate-grown Methanosarcina acetivorans. BMC Microbiol. 11, 165. doi: 10.1186/1471-2180-1111-1165 (2011).

