## Supplemental for "Carbonic anhydrase plays multiple roles in acetotrophic growth of a model marine methanogen from the domain *Archaea*"

**This PDF file includes:**

Figure S1. Sequence alignment of Cam from *M. acetivorans* and *M. thermophila*.
Figure S2. Transmembrane topology and signal peptide prediction (https://phobius.sbc.su.se/).
Figure S3. Sequence alignment of CamH from *M. acetivorans* and *M. thermophila*.
Figure S4. Sequence alignment of CamH and Cam from *M. acetivorans*.
Table S1. Methane production in resting cell suspensions containing acetate or methanol.

```
M. thermophila P40881 MMFNKQIFTILILSLSLALAGSGCISEGAEDNV---AQEITVDEFSNIRENPVTPWNPEP 57
M. acetivorans Q8TMW3 -MKINRIFLALLFSLALTLAGSGCVSQGEGAEDGESADTEVESEVSNIRANPVTPWNPEP 59
                       *  ::**  *::**:*:******:*:*   :    *:  . .*.**** **********
                                                                         %
M. thermophila P40881 SAPVIDPTAYIDPQASVIGEVTIGANVMVSPMASIRSDEGMPIFVGDRSNVQDGVVLHAL 117
M. acetivorans Q8TMW3 TEPVIDSTAYIHPQAAVIGDVTIGASVMVSPMASVRSDEGTPIFVGDETNIQDGVVLHAL 119
                      : **** ****.***:***:*****.********:***** ******.:*:*********

                      +                                %    %
M. thermophila P40881 ETINEEGEPIEDNIVEVDGKEYAVYIGNNVSLAHQSQVHGPAAVGDDTFIGMQAFVFKSK 177
M. acetivorans Q8TMW3 ETVNEEGEPVESNLVEVDGEKYAVYVGERVSLAHQSQIHGPAYVGNDTFIGMQALVFKAN 179
                      **:******:*.*:*****::****:*:.********:**** **:********:***::

                                                                          +
M. thermophila P40881 VGNNCVLEPRSAAIGVTIPDGRYIPAGMVVTSQAEADKLPEVTDDYAYSHTNEAVVYVNV 237
M. acetivorans Q8TMW3 VGDNCVLEPKSGAIGVTIPDGRYIPAGTVVTSQAEADELPEVTDDYGYKHTNEAVVYVNV 239
                      **:******:*.*************** *********:********.*.***********

M. thermophila P40881 HLAEGYKETS 247
M. acetivorans Q8TMW3 NLAAGYNA-- 247
```

**Figure S1. Sequence alignment of Cam from *M. acetivorans* and *M. thermophila*.** Symbols identify catalytically relevant residues (+) and metal ligands (%) for *M. thermophila*. Acidic loop residues, including the proton shuttle residue Glu84, are underlined. Shaded residues indicate the post translationally cleaved leader peptide. Sequences were obtained from UniProt (www.uniprot.org). Alignment was with CLUSTAL O (1.2.4).

```
ID   Cam
FT   SIGNAL         1      26
FT   REGION         1       6       N-REGION.
FT   REGION         7      18       H-REGION.
FT   REGION        19      26       C-REGION.
FT   TOPO_DOM      27     247       NON CYTOPLASMIC.
//
```

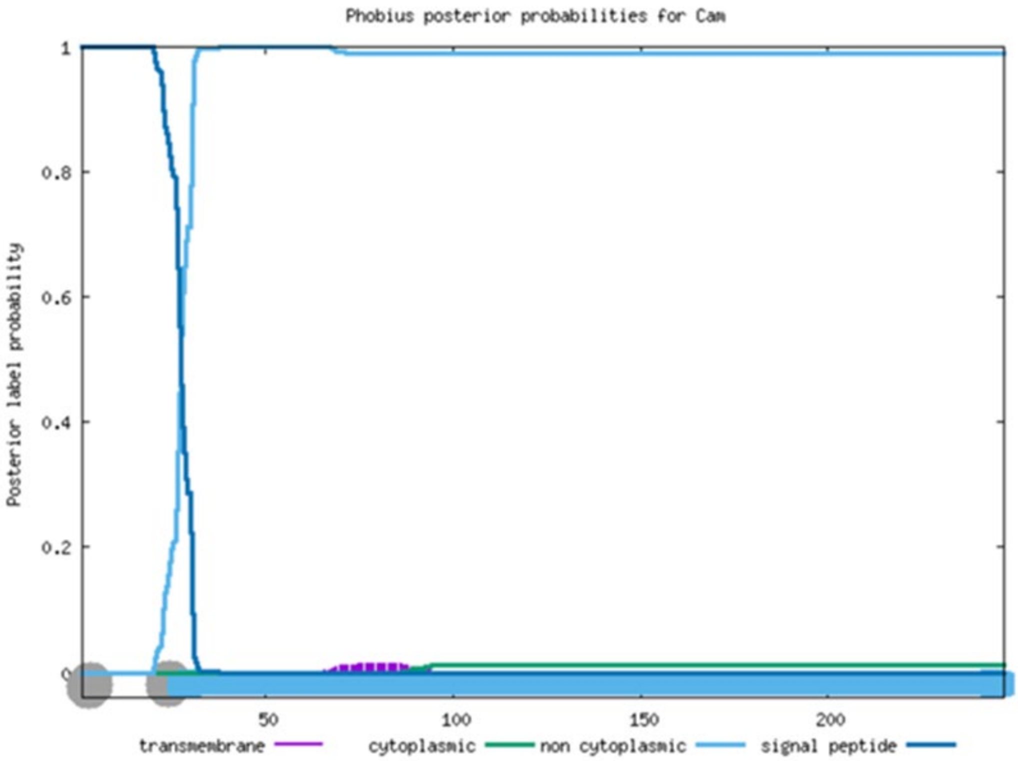


**Figure S2. Transmembrane topology and signal peptide prediction**

**(https://phobius.sbc.su.se/).**

```
M. thermophila C3W4Q7 MKRNFKMHLPNPHKQHPKVSKRAWISETALIIGNVSIADDVFVGPNAVLRADEPGSSITV 60
M. acetivorans Q8TRS1 ------MNLPNPRKQHPRVSKRAWISETAVIIGNISIADYVFVGPNAVLRADEPGSSITV 54
                            *:****:****:***********:****:**** ********************

M. thermophila C3W4Q7 HRGCNVQDNVVVHSLSHSEVLIGKNTSLAHSCIVHGPCRIGEDCFIGFGAVVFDCNIGKD 120
M. acetivorans Q8TRS1 QSGCNVQDNVVVHSLSHSDVLVGKNTSLAHSCIVHGPCRIGEGCFIGFGAVVFDCNIGKD 114
                      : ****************:**:********************.*****************

M. thermophila C3W4Q7 TLVLHKSIVRGVDISSGRMVPDGTVITRQDCADALEDITKDLTEFKRSVVKANIDLVEGY 180
M. acetivorans Q8TRS1 TLVLHRSVVRGIDIFSGRIVPDGTVITRQAYANALEPITKEMTEFKRSVVRANIELVEGY 174
                      *****:*:***:** ***:**********  *:*** ***::********:***:*****

M. thermophila C3W4Q7 IRLREES 187
M. acetivorans Q8TRS1 MKLREES 181
                   ::*****
```

**Figure S3. Sequence alignment of CamH from *M. acetivorans* and *M. thermophila*.**

Sequences were obtained from UniProt (www.uniprot.org). Alignment was with CLUSTAL O (1.2.4).

```
M. acetivorans CamH Q8TRS1 ---------------------------------------------------MNLPNPRKQ 9
M. acetivorans Cam  Q8TMW3 MKINRIFLALLFSLALTLAGSGCVSQGEGAEDGESADTEVESEVSNIRANPVTPWNPEPT 60
                                                                              :.  **.

M. acetivorans CamH Q8TRS1 HPRVSKRAWISETAVIIGNISIADYVFVGPNAVLRADEPGSSITVQSGCNVQDNVVVHSL 69
M. acetivorans Cam  Q8TMW3 EPVIDSTAYIHPQAAVIGDVTIGASVMVSPMASVRSDE-GTPIFVGDETNIQDGVVLHAL 119
                           .* :.. *:*   *.:**:::*.  *:*.* * :*:** *: * * .  *:**.**:*:*

M. acetivorans CamH Q8TRS1 S-------------------HSDVLVGKNTSLAHSCIVHGPCRIGEGCFIGFGAVVFDCN 110
M. acetivorans Cam  Q8TMW3 ETVNEEGEPVESNLVEVDGEKYAVYVGERVSLAHQSQIHGPAYVGNDTFIGMQALVFKAN 179
                           .                   :  * **:..****.. :***. :*:. ***: *:**..*

M. acetivorans CamH Q8TRS1 IGKDTLVLHRSVVRGIDIFSGRIVPDGTVITRQAYANALEPITKEMT--EFKRSVVRANI 168
M. acetivorans Cam  Q8TMW3 VGDNCVLEPKSGAIGVTIPDGRYIPAGTVVTSQAEADELPEVTDDYGYKHTNEAVVYVNV 239
                           :*.: ::  :* . *: * .** :* ***:* ** *: *  :*.:    . :.:** .*:

M. acetivorans CamH ELVEGYMKLREES 181
M. acetivorans Cam  NLAAGYNA----- 247
                    :*: **
```

**Figure S4. Sequence alignment of CamH and Cam from *M. acetivorans*.** Sequences were obtained from UniProt (www.uniprot.org). Alignment was with CLUSTAL O (1.2.4).

| Experimental condition | | | Rates of methane production (nmole/min x $mg^{-1}$ protein) | | |
|---|---|---|---|---|---|
| **Strain** | **0.15 atm $CO_2$** | **substrate** | **100 mM (500 µmol)** | **20 mM (100 µmol)** | **5 mM (25 µmol)** |
| wild-type | + | acetate | 58.4 ± 1.6 | 46.9 ± 2.9 | 42 ± 3.6 |
| Δ*cam* | + | acetate | 40.2 ± 1.8 | 9.1 ± 0.5 | none detected |
| wild-type | - | acetate | none detected | none detected | none detected |
| Δ*cam* | - | acetate | none detected | none detected | none detected |
| wild-type | + | methanol | 60.4 ± 1.6 | not determined | 61.3 ± 2.2 |
| Δ*cam* | + | methanol | 59.4 ± 2.2 | not determined | 59.8 ± 2.1 |
| wild-type | - | methanol | 61.5 ± 1.7 | not determined | 62.4 ± 1.8 |
| Δ*cam* | - | methanol | 61.8 ± 1.8 | not determined | 61.2 ± 1.7 |

**Table S1. Methane production in resting cell suspensions containing acetate or methanol.** Cell material was from cultures grown with the indicated substrate. Values are amounts of methane produced after addition of substrate. Cells containing 5 mg protein were used in each experiment. Values are the mean of four biological replicates with standard deviation. No methane was detected in controls without added substrate.
